# A survey to assess animal methods bias in scientific publishing

**DOI:** 10.1101/2022.03.24.485684

**Authors:** Catharine E. Krebs, Ann Lam, Janine McCarthy, Helder Constantino, Kristie Sullivan

**Author notes:** **Corresponding author** Catharine E. Krebs, PhD, Physicians Committee for Responsible Medicine, 5100 Wisconsin Ave., NW, Suite 400, Washington, DC 20016-4131.

## Abstract

Publication of scientific findings is fundamental for research, pushing innovation and generating interventions that benefit society, but it is not without biases. Publication bias is generally recognized as journal’s preference for publishing studies based on the direction and magnitude of results. However, early evidence of a newly recognized type of publication bias has emerged in which journal policy, peer reviewers, or editors request that animal data be provided to validate studies produced using nonanimal-based approaches. We describe herein “animal methods bias” in publishing: a preference for animal-based methods where they may not be necessary or where nonanimal-based methods may be suitable, which affects the likelihood of a manuscript being accepted for publication. To gather evidence of animal methods bias, we set out to collect the experiences and perceptions of scientists and reviewers related to animal- and nonanimal-based experiments during peer review. We created a survey with 33 questions that was completed by 90 respondents working in various biological fields. Twenty-one survey respondents indicated that they have carried out animal-based experiments for the sole purpose of anticipating reviewer requests. Thirty-one survey respondents indicated that they have been asked by peer reviewers to add animal experimental data to their nonanimal study; 14 of these felt the request was sometimes justified, and 11 did not think it was justified. The data presented provide preliminary evidence of animal methods bias and indicate that *status quo* and conservatism biases may explain such attitudes by peer reviewers and editors.

## Introduction

Publication of scientific results is a necessary step in the dissemination and implementation of biomedical advances and plays a central role in the progress of researchers’ careers. It is a fundamental part of the research process that promotes knowledge sharing, pushes innovation, and contributes to the development of research standards and interventions that benefit society. However, scientific publication in the biological and biomedical fields has been greatly affected by practices such as publication bias, which is currently most recognized as a journal’s preference for publishing studies based on the direction and magnitude of results, while disfavoring those without statistical significance (1).

More recently, early evidence of what may be a new type of publication bias in biomedical and health sciences has emerged. This type of bias occurs when the likelihood of a study being published is affected by the methods used. For instance, in 2018, a preprint of a study using human airway epithelial organoids and resected tissue from patients with cystic fibrosis used no animal-based experiments (2). However, in the accepted version of the same study published in the EMBO journal, additional animal-based experiments were presented in which tumor airway-organoids were transplanted into immunocompromised mice (3). According to Hans Clevers, the senior author of the study and research group leader at Utrecht University in the Netherlands, *de novo* animal data was produced and added to the new version of the study manuscript because of requests by peer reviewers (4).

Since then, a perspective article was written by Donald Ingber at the Wyss Institute asks: “Is it Time for Reviewer 3 to Request Human Organ Chip Experiments Instead of Animal Validation Studies?” (5). In this article, the author questions why animal data is still considered the gold standard by reviewers in biomedical research while presenting evidence that organ chips may better suit this purpose. This question reinforces the anecdote of the airway organoids study described above, revealing a publication bias caused by a status quo reliance on animal-based methods. That is, bias created by editorial policy, peer reviewers, or editors insisting on the inclusion of animal-based evidence, sometimes as a condition for acceptance of the manuscript. In this paper, we examine this phenomenon, which we term “animal methods bias.” Specifically, animal methods bias indicates a reliance on or preference for animal-based methods despite the availability of potentially suitable nonanimal-based methods. Animal methods bias may occur during the review of grant applications as well, but the current study focuses on animal methods bias in publishing.

Nonanimal-based methods have advanced a great deal in the past decade, improving on the limitations of animal-based methods with their ability to replicate the molecular, cellular, and physiological mechanisms of human diseases more reliably and to translate preclinical findings into safe and effective treatments for patients in a more predictive manner. Human organoids, or stem cell-derived 3D culture systems, can replicate the structure and function of human organs (6) and have been used to model many human organ systems including brain (7), lung (8), and intestines (9). Organoids have been used in a variety of applications such as investigating infectious diseases, genetic disorders, and cancer, and to enable precision medicine (6). Similarly, organ chips are composed of human stem or primary cells, but cells are situated in microfluidic systems allowing fluid flow to mimic dynamic processes like circulation and tissue-tissue interfaces (5).

Nonanimal-based methods like organoids and organ chips are powerful tools that can replace the use of animals in many applications, but there remain many barriers to their uptake, including biases like the one described here. Thus, animal methods bias may have far-reaching consequences, including (i) the continued use of animal-based research methods despite their unreliability in modeling diseases and predicting success in clinical trials, (ii) stifled use and further development of nonanimal methods, despite purported commitments to the principles of the 3Rs, and (iii) misdirected prioritization of federal research dollars. There is therefore a scientific and moral imperative to address this bias and mitigate its consequences for researchers and patients.

Although anecdotal accounts of animal methods bias have been described, there is not yet to our knowledge a published study that describes or presents evidence of animal methods bias in publishing. To begin investigating this issue systematically, we designed a survey study. The aim was to understand the experiences and perceptions of scientists and reviewers related to animal methods bias in publishing. To this end, we created a survey and shared it through social media and our private channels. This paper presents preliminary evidence of animal methods bias in publishing, discusses the many implications of this bias, and presents potential solutions to address it.

## Material and Methods

### Survey questions and logic

Survey questions were designed to collect the experiences and perceptions of researchers and peer reviewers on the topic of animal methods bias in publishing peer review (Supplementary Material: S1 Appendix). Participants were asked up to 33 questions including up to 25 multiple choice and up to eight open-ended questions. Decision logic was used to route respondents through the survey based on their answers, thereby resulting in a varying number of respondents answering each question (Supplementary Material: S2 Appendix). For example, because we were only interested in experiences during peer review, respondents who indicated that they had zero peer reviewed publications were routed to the end of the survey.

### Survey dissemination

The survey was designed and collected using SurveyMonkey. A short introduction to the survey was provided to respondents before the survey questions were presented, which included a brief statement of purpose, as well as a confidentiality statement:

“This survey aims to better understand the circumstances under which animal-derived data is requested to be added to a study performed without the use of animals as a condition for publication. The questions of this survey were designed to collect the experiences and perceptions of both researchers and reviewers on this topic. We thank you in advance for your participation and encourage you to disseminate this survey to your colleagues. Your responses will remain anonymous, except to surveyors in the case that you choose to provide your name and contact details, which will remain confidential. Responses will not be used for a discriminatory purpose.”

Because survey respondents may consider animal and nonanimal experiments to have an array of different meanings, and to avoid any confusion on how to classify types of study, we defined the two concepts in the introduction of the survey:

Animal-based experiment: An experiment performed in a living non-human animal or in a non-human animal-derived organ, tissue, or other biological product, e.g., an animal *in vivo* model or animal cell-derived in vitro model.

Nonanimal-based experiment: An experiment performed in a living human or in a human-derived organ, tissue, or other biological product, or in a nonanimal specimen, or in silico, e.g., human cell-derived in vitro model, even if it uses animal-based materials such as buffers or antibodies, or a purely computational model.

A URL link to the survey was sent out by members of the research team to LinkedIn groups related to life sciences, to colleagues at other institutions and organizations, some of whom shared the survey link through official organizational communications, and via research team organizational communications. The survey was open to responses from May 20, 2021 until August 14, 2021. The close date was decided in advance to ensure that no survey responses would be collected after a presentation on the survey was given at the World Congress on Alternatives and Animal Use in the Life Sciences (https://www.wc11maastricht.org/) conference on August 31, 2021.

### Analysis

Survey data were downloaded from SurveyMonkey following the close of the survey and imported into Excel and R studio for processing and analysis (Supplementary Material: S3 Data). Due to the small sample size, survey questions were analyzed in descriptive and qualitative manners only by taking counts and percentages but not performing statistical tests. As such, multiple choice answers to questions about the frequency of experiences during manuscript review were primarily considered as binary, assessing whether something had occurred never or ever. In addition, the survey results reflect a qualitative assessment of whether the animal methods bias phenomenon occurs, not to what extent it occurs. When reporting on open-ended prompts (Boxes 1-6), we edited the answers for clarity. Edits are indicated with brackets: [].

## Results

### Respondent demographics

There was a total of 90 respondents. Nine respondents indicated that they had zero peer reviewed publications and an additional 13 respondents failed to complete survey questions about their research and experiences in publishing peer review. These respondents were excluded from analysis, resulting in a sample size of 68 respondents (Tab. 1).

**Table 1.** Respondent demographics.

| Characteristic | N | Percent total |
| --- | --- | --- |
| *Total* | 68 |  |
| Sector (question 2) |  |  |
| Academia/Research | 50 | 74% |
| Industry | 7 | 10% |
| Government | 4 | 6% |
| Nonprofit/NGO | 3 | 4% |
| Publishing | 1 | 1% |
| Gender (question 3) |  |  |
| Male | 35 | 51% |
| Female | 31 | 46% |
| Nonbinary | 1 | 1% |
| Prefer not to say | 1 | 1% |
| Highest degree earned (question 5) |  |  |
| Master's degree | 5 | 7% |
| Research doctorate | 52 | 76% |
| Medical degree | 20 | 29% |
| Years since highest earned degree (question 6) |  |  |
| 0-5 | 11 | 16% |
| 6-10 | 19 | 28% |
| 11-20 | 14 | 21% |
| >20 | 24 | 35% |
| Number of peer-reviewed publications (question 7) |  |  |
| 1-10 | 23 | 34% |
| 11-20 | 9 | 13% |
| 21-40 | 11 | 16% |
| 41-100 | 14 | 21% |
| >100 | 11 | 16% |

There was a broad distribution across research fields. The fields that were most represented were medicine and clinical research (28%), molecular and cellular biology (21%), neuroscience (15%), pharmacology (16%), and toxicology (19%; Tab. S1). The geographic representation of respondents was not evenly distributed. The largest number of respondents primarily worked in the United States (32%), followed by South Korea (15%; Tab. S1). Most respondents (75%) worked in academia or for a research institution (Tab. 1). There was a slightly greater number of male respondents (51%; Tab. 1). Most respondents held a research doctorate as their highest earned degree (76%; Tab. 1). To assess the level of seniority and extent of experience in peer review, respondents were asked how many years it had been since they received their highest earned degree and number of peer-reviewed publications, both of which showed fairly even distributions (Tab. 1).

### Respondent use of animal and nonanimal methods

To help understand respondents’ internal biases related to the nature of research methods they regularly use, respondents were asked about animal and nonanimal methods used in their research. Respondents’ answers were slightly skewed toward using nonanimal research models, samples, data, or materials (68%) versus animal research models, samples, data, or materials (56%; Tab. 2). Asked another way, 44% of respondents indicated that they never use animal experiments, while 56% indicated that they use animals at least rarely (Tab. 2).

**Table 2.** Respondent use of animal and nonanimal methods in research. For types of models, sample, data, or materials (survey question 8), respondents could choose as many answers as applicable.

|  | N | Percent<br>Total |
| --- | --- | --- |
| *Total* | 68 |  |
| Type of models, samples, data, or materials<br>(question 8) |  |  |
| Animal | 38 | 56% |
| Nonanimal | 46 | 68% |
| Frequency of animal use (question 10) |  |  |
| Never | 30 | 44% |
| Rarely | 13 | 19% |
| Sometimes | 14 | 21% |
| Often | 4 | 6% |
| Always | 7 | 10% |
| Use of animals for sole purpose of anticipating<br>reviewer requests (question 11) |  |  |
| Never | 47 | 69% |
| Rarely | 8 | 12% |
| Sometimes | 9 | 13% |
| Often | 4 | 6% |
| Always | 0 | 0% |

These data indicate a possible bias in the perceptions of respondents’ experiences related to requests for animal experiments during manuscript review. When asked how often respondents use animal experiments for the sole purpose of anticipating reviewer requests for them, most respondents (69%) indicated that they never do this (Tab. 2). However, eight respondents indicated that they do this rarely, nine do this sometimes, and four do this often, indicating that journals’ perceived reliance on animal evidence may contribute to the performance of potentially unnecessary experiments.

### Respondent experiences as authors during manuscript peer review

To assess researcher experiences during manuscript peer review regarding requests for animal experiments, survey prompts were provided regarding the request by reviewers to add animal experimental data to their nonanimal study. The sample size (n=68) was too small to determine the extent to which this occurs. As such we consider the answers to these questions as indicators of the presence of animal methods bias rather than a quantification of the frequency of occurrence or of the proportion of researchers to whom this happens. Thirty-two respondents indicated that they have never been asked for inclusion of animal data, whereas 31 respondents indicated that this has happened at least one time (Fig. 1).

**Figure 1.**
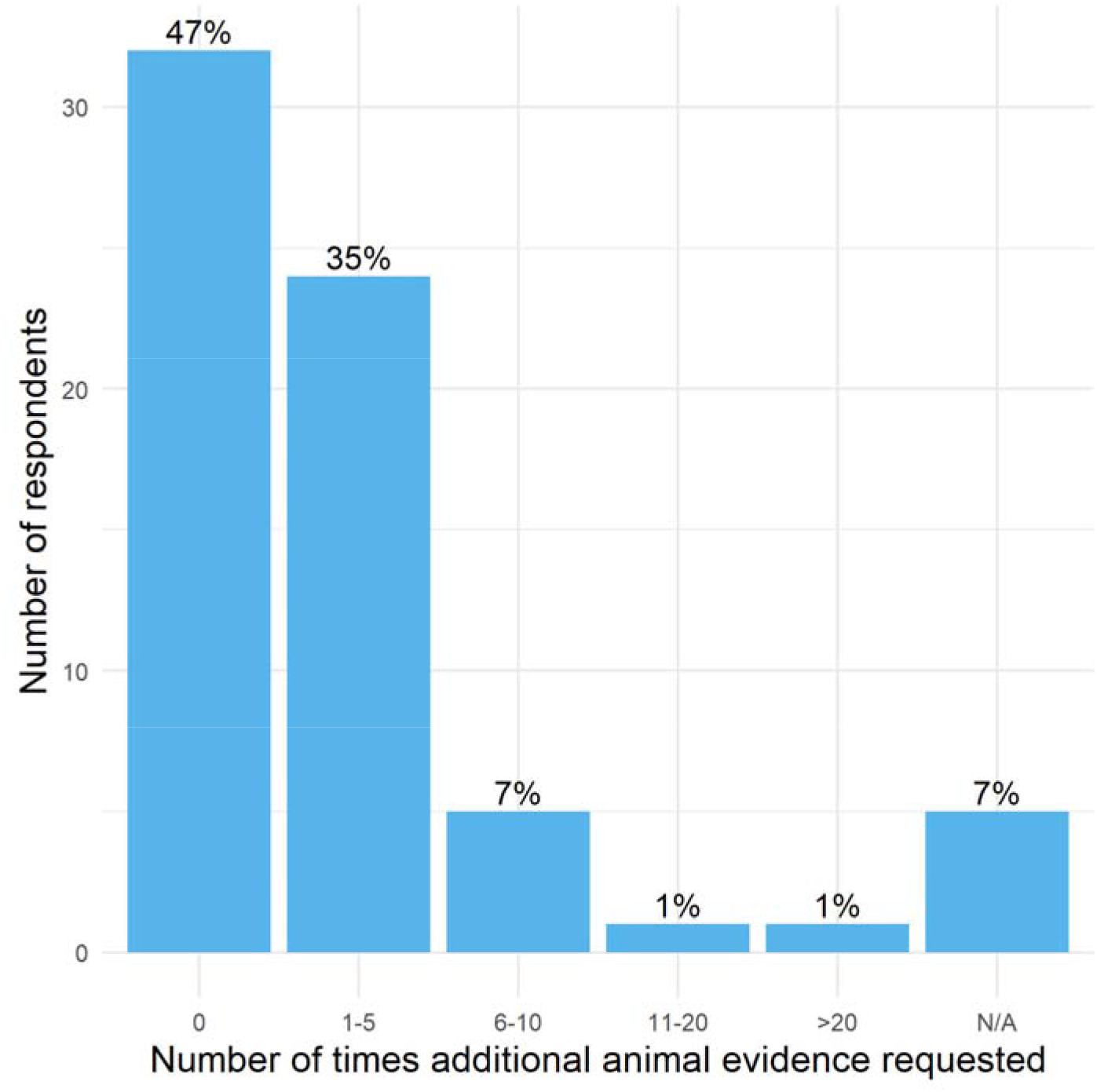
Number of times respondents have been requested by reviewers for animal data for a nonanimal study. Responses to question 12: “During manuscript submission peer review, how many times have you been asked for animal experimental data to be added to a study that otherwise had no animal-based experiments?” Thirty-two respondents indicated that they have never been asked for inclusion of animal data; 31 respondents indicated that this has happened at least one time. The percent of respondents answering in each category is on top of each bar (total N=68).

Of the 31 respondents who indicated that additional animal-based data was requested, only three respondents thought that the request was justified, 14 respondents felt that it was sometimes justified, and 11 respondents did not think the request was justified. Three respondents did not provide an answer to this question. When asked to elaborate on their answers in an open-ended prompt, further information was gained about researcher experiences (Box 1). These respondents were then asked if they complied with the request for additional animal-based experiments to be added to a study that otherwise had no animal experiments and to elaborate on their answers, to which 14 indicated that they did not comply, nine indicated that they sometimes complied, and five indicated that they did comply (Box 2).

#### Box 1.

**Respondents’ feelings toward the justification of requests for additional animal experimental evidence**

Responses to question 14: open-ended elaborations to question 13, “Did you feel the requested additional animal-based experiments were justified?”; grouped by whether respondents felt it was justified.

##### Not justified

Animal models are seen as a way to validate *in vitro* data. This is [ironic].

The study was about heterogeneity of cancer cells from human tissue samples. It was irrelevant to do an experiment on mice.

The reviewer simply claimed that the human cellular models we used were not useful models to study viral infection, implying that only animal models would be acceptable.

Referees ask for animal experiments because it is customary to do so in the field of biophysics, toxicology not because it is necessary. Many researchers are unaware about the potential of *in vitro, in silico* methods and human based models.

The animal [studies] that were asked were to only [emphasize] the results we obtained *in vitro*. [N]o other reason.

[M]erely requested to confirm *in vitro* data, no added value.

The need for validation of human organoid data with animal studies… just because journal reviewers were used to this.

They wanted human *in vitro* data to be “validated” against an animal model.

We elucidated a [mechanism of action] using only gene expression data from hepatocytes *in vitro*. We were asked to perform the same *in vivo*, to confirm the findings.

##### Sometimes justified

Some [evidence is] lacking with *in vitro* experiments (lack of previous research data linking to clinical data).

It was primarily a reasonable request to verify the efficacy of the substance. However, there were some cases where the importance was not very high for the completeness of the thesis.

IRB

It is not justified to ask [a] modeler, human experimenter to replicate their findings in [an] animal [experiment], because I have no lab to do this [and] present plausible model simulations that support my case. I would need to set up a collaboration to get these animal data, [and that] would take additional years, in the meantime papers are not published and others cannot build upon the work.

To understand interaction [between] a tumor cell and host immune system or host cells in general or to study systemic reactions to a drug animal experiments are justified in my opinion.

Sometimes the reviewers identify critical gaps in knowledge, these are valuable peer reviews. Other times it seems like they ask for animals out of habit. We refuse. This is even more difficult and hard to deal with when it comes to grant reviews

Animal based models are not always the best in specific research. In cancer research some level of animal data is sometimes required.

In some circumstances the added information would indeed provide extra validation, but it was not essential.

##### Justified

Some ethical and social limitations can be overcome and long-term complete observation can be made.

#### Box 2.

**Respondents’ thoughts on complying with requests for additional animal experimental evidence**

Responses to question 16: open-ended elaborations to question 15, “Did you comply with the request(s)?”; grouped by whether respondents complied.

##### Did not comply

With limited time/money along with ethical concern, request was rejected and paper was withdrawn.

We submit[ed] in a different journal.

Reviewer three said words to the effect of “there are no mouse experiments in this study therefore I reject it for this journal.” They said nothing about the rest of the study. I submitted to a different journal.

We submitted the manuscript to another journal.

Submitted to another journal as we [don’t] have an animal breeding facility as we comply with 3R.

My group is purely computational.

We wrote a rebuttal to the editor, no success. Submitted elsewhere (high impact) accepted within 2 months.

Justified why I shouldn’t have to.

The paper was withdrawn and used only as chapter in my PhD thesis.

##### Sometimes complied

I have responded to reasonable requests.

I have pulled publications and waited for [animal] labs to perform [experiments] with my stimuli first. I gave them the support to run these [experiments], but [I] am not necessarily an author on that work, and this causes massive delays in the publications from my lab.

In my early days of research [I] did not [have] access to animal models therefore I was able to comply with the requests only when establishing dedicated collaborations with [colleagues].

[Sometimes animal experiments] were just not applicable.

In my group sometimes animal experiments are performed to publish [in journals] with high impact [factors] but I do not agree. I would go on employing and studying new non-animal methods.

[To] satisfy reviewers’ request flank (subcutaneous) injections on tumor cells were performed.

##### Complied

In [one] study I can think of we performed mouse xenograft experiments

Respondents who indicated that they complied with the request(s) were then asked who performed the additional animal experiments, to which five indicated that their own lab(s) or lab(s) of coauthors did, two indicated that additional collaborators were brought in, and five indicated that some combination of their own labs and additional collaborators performed the experiments. When asked to elaborate on these answers, some respondents indicated that coauthors or members of their own lab had access to animal facilities and one respondent indicated that a lab member had “experience with performing *in vivo* procedure” (Box 3).

#### Box 3.

**Respondents’ elaborations on when complying with the request, who performed the additional experiments**

Responses to question 18: open-ended elaborations to question 17, “When you complied with the request(s), did members of your own lab or the labs of coauthors perform additional experiments, or did you and coauthors bring in additional collaborators?”; grouped by who performed the additional experiments.

##### Own lab(s) or lab(s) of coauthors

We have IACUCs in limited circumstances. If we need to do the work, we do it - although replacement is important, we’re not at the level of 100% replacement yet.

A coauthor had this facility.

A member of my lab had access to animal facility and experience with performing the *in vivo* procedure.

##### Additional collaborators

I do not have an animal lab.

##### Combination

Sometimes they are performed by the coauthors.

Respondents who indicated that they did not comply with the request(s) for additional animal experimental evidence were then asked if they have ever had a manuscript rejected because they did not comply. Nine respondents indicated that they have not had a manuscript rejected for not complying with a request for animal experiments and eight indicated that they weren’t sure, but nine indicated that they have. Elaborations on these answers are provided in Box 4. Finally, respondents were asked in an open-ended prompt to provide any additional details about their experiences being asked for additional evidence during manuscript review (Box 5).

#### Box 4.

**Respondents’ thoughts on having a manuscript rejected for not complying with a request for additional animal experiments**

Responses to question 20: open-ended elaborations to question 19, “Have you ever had a manuscript rejected because you did not comply with a request for animal experimental data to be added to a study that otherwise had no animal-based experiments?”; grouped by whether this has happened to them.

##### No

I just submitted to other journals if animal studies were requested. Cannot remember a case with over 400 publications.

[Normally], when *in vivo* data were provided, articles were accepted.

##### Not sure

Usually, that’s not only the reason.

If there [were] requests, we accepted all requests and did the experiments.

I know from the experience of my PI that papers not showing animal experiments are much more likely to be rejected from high impact journals.

##### Sometimes

[Some evidence is] lacking with *in vitro* experiments (lack of previous research data linking to clinical data).

Sometimes the reviewers identify critical gaps in knowledge, these are valuable peer reviews. Other times it seems like they ask. for animals out of habit. We refuse. This is even more difficult and hard to deal with when it comes to grant reviews

##### Yes

They did not trust the *in silico* predictions in [a biophysical] realistic computational model, and wanted the animal evidence.

Yes. [The] reviewer said [we were] missing animal results. When [I] sent my response back that [I won’t] do it, it was rejected.

#### Box 5.

**Additional details respondents shared about being asked for additional experimental data**

Responses to question 24: “Are there any additional details you would like to share about your experience(s) during manuscript review being asked for additional experimental data?”

Personally, I think it’s [a] rare [case] to ask to add unreasonable or unnecessary animal data.

In my experience, reviewers on grant panels are even more likely to ask for animal experiments than manuscript reviewers.

The journals that wanted animal experiments were journals with higher impact [factors]. I submitted my work to journals with lower impact [factors]. Therefore, it is not [too] hard to conclude that people who do animal experiments are perceived to be better scientists because they are allowed/encouraged to submit to those journals with higher impact [factors]. [It is a self]-fulfilling prophecy, meaning there is no incentive to not do animal experiments. One scientist at a conference proudly announced they had used 2,000 mice in one paper. I went to another seminar where the whole talk was about a virus study in primates; at the end they showed that the effect in primates was totally different to that in humans, therefore negating their entire study.

I don’t remember being asked to perform experiments on animal models when I was not the principal investigator of the projects, but it is possible that the principal investigator decided to do experiments on animals just because it was the normal thing to do.

I have been asked several times to peer review animal-based studies. I have always declined. I work in perinatal neurology and obtain my data mainly from imaging (MRI, ultrasound). I have seen animal-based studies where a brain injury was caused to an animal (during gestation or first few postnatal days) with the aim to show the pattern of injury or developmental consequences. Most of the data are already known from human studies. I have always written to editors explaining why they should not accept this kind of manuscripts; I have never received a response. The number of manuscripts submitted with these animal data is still huge.

### Respondent experiences as reviewers during manuscript peer review

We then asked a series of questions aimed at gaining insight into the experiences of reviewers requesting the addition of animal experimental evidence. Of the 57 respondents remaining (several had dropped out of the survey by this point), six indicated that they never serve as a manuscript reviewer for scientific journals, and the remaining 51 were routed to a series of questions specifically for reviewers (Fig. S1). These respondents were then asked, in their capacity as a reviewer for a scientific journal, how often they requested animal experimental data to be added to a study that otherwise had no animal-based experiments. Of these 51 respondents, 36 said they never made these requests, but 15 indicated that they did (Fig. 2).

**Figure 2.**
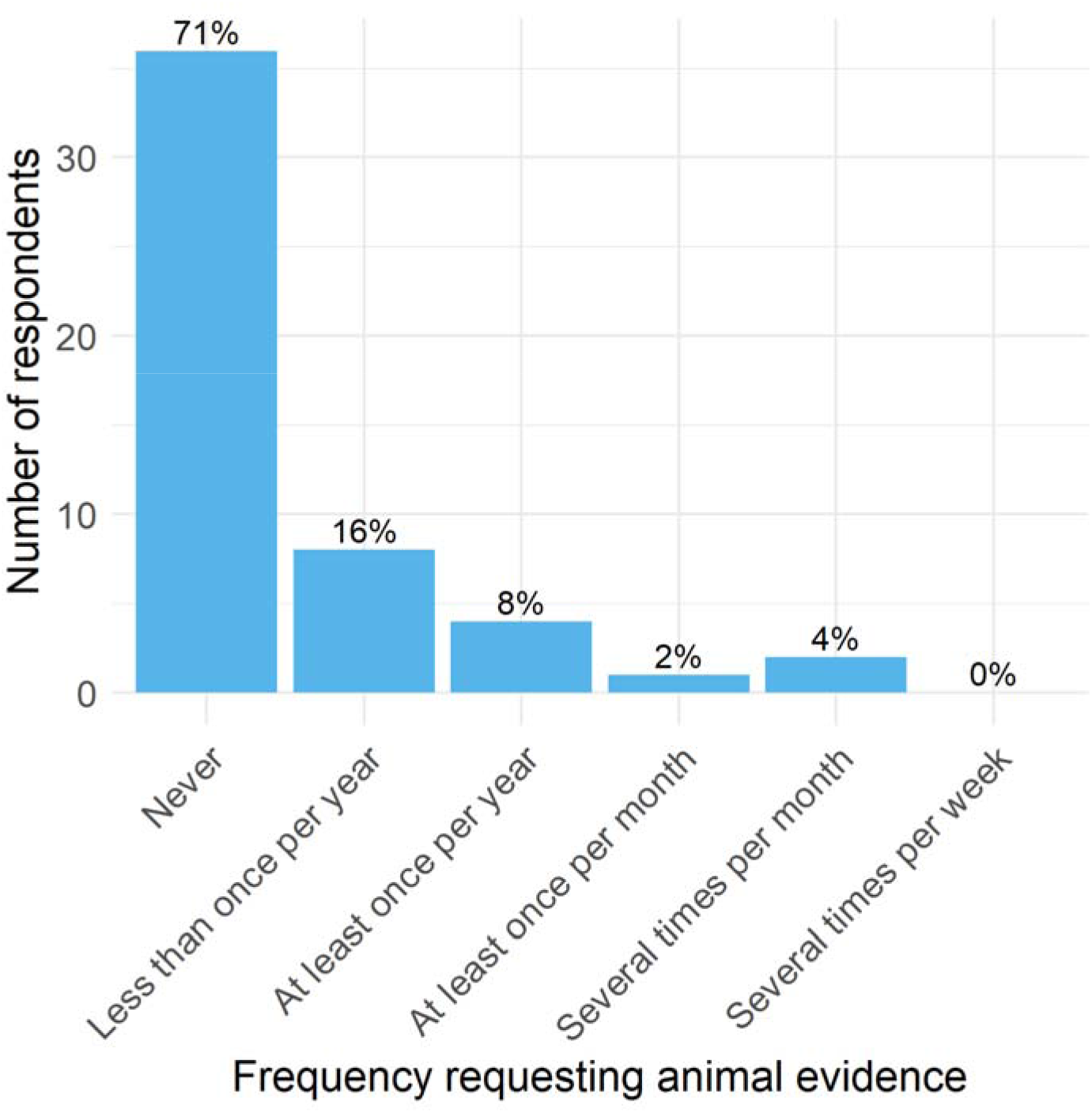
Respondent frequency of requesting as a reviewer animal data for a nonanimal study. Responses to question 26: “In your capacity as a reviewer for a scientific journal, how often do you request animal experimental data to be added to a study that otherwise had no animal-based experiments?” Thirty-six respondents indicated that they never made these requests; 15 indicated that they did less than once per year or more. The percent of respondents answering in each category is on top of each bar (total N=51).

We then asked these 15 respondents for what reasons they made these requests, to which one responded due to a “Request from editor,” six responded due to “Preference for animal-based methods,” and seven responded because they are “Unaware of nonanimal or alternative methods for the given hypothesis.” Respondents were asked to elaborate on the previous answer (Box 6). Finally, respondents had the opportunity to provide any additional thoughts as reviewers they wanted to share (Box 7).

#### Box 6.

**Respondent elaborations on reasons for requesting additional animal-based experimental data**

Responses to question 28, open-ended elaborations to question 27, “For what reasons have you requested that animal-based experimental data be added to a study that otherwise had no animal experiments?”

When I think that it is difficult to verify a hypothesis with only *in vitro* experiments, I request *in vivo* verification for systemic evaluation.

Sometimes, the interaction between host and pathogens cannot be modelled at organism levels.

Non animal models don’t represent actual biology and kinetics

#### Box 7.

**Additional details respondents shared about their experience as a reviewer requesting additional data**

Responses to question 32: “Are there any additional details you would like to share about your experience(s) as a reviewer requesting additional data?”

Since animal experiment proceeds with a lot of consideration, it is rare that indiscriminate additional experiments are required.

The type of extra experiments I request depends mainly on the focus of the study and the hypothesis the authors want to demonstrate. Therefore, I suggest animal experiments more rarely than *in vitro* experiments. This also depends on the knowledge that often extra animal experiments require ethics approval and a longer time to be performed therefore they are not always compatible with the timings for resubmission allowed by the publishers. If a substantial amount of animal experiments is requested, in my opinion, I [would] rather suggest rejection than revision.

My main request for additional data is when controls are missing - this seems to be the most common missing thing these days - I would say it is more of a problem than it was ∼20 years ago as people rush to publish.

In some cases the original animal experiment was not conducted properly or without the right controls. Then I would make suggestions.

If I am asked to review a paper that is entirely based on mouse models for example, I will probably not accept to review this because I don’t work with mouse models myself. If there are lots of other molecular and cellular experiments with a few mouse experiments, then I would accept that paper to review.

Often researchers are not aware of newly developed methods. As referee it is important to share info.

I still didn’t have to ask for additional experiments to improve a manuscript, I don’t have that much experience being both principal investigator or reviewer. I don’t like animal experiments and consider most of them unnecessary, so it is very unlikely that I would request additional experiments with animals for a publication.

The type of data I have requested is better or additional statistical analysis.

I get data since rationale is there. Reproducibility and real time data is required for drugs.

Sometimes additional experiments are needed to gather further mechanistic understanding of described results, and nonanimal experiments serve this purpose.

## Discussion

In this study, we set out to investigate animal methods bias in publishing through a survey initially completed by 90 respondents working in several areas in the biological and biomedical fields. The data presented provide preliminary evidence of animal methods bias. Here we discuss the potential causes and implications of this type of bias and possible solutions to address it.

To our knowledge, there was no prior specific method or strategy to investigate the occurrence and frequency of animal methods bias in publishing. Thus, using the SurveyMonkey platform, we designed a survey called “Survey to Assess Journal and Reviewer Requests for Evidence in Animals.” Based on our survey, we cannot tell how frequently this type of bias occurs because our sample size of 90 was too small to conduct statistical tests. Furthermore, it is possible that our dissemination strategy may have led to a nonrepresentative sample comprised of researchers who would like to avoid using animals in research. Indeed, the survey was partially disseminated through our professional networks and social media channels, which predominantly include researchers who preferentially or exclusively use nonanimal methods. Thus, a larger survey that is sufficiently powered and representatively sampled will be necessary to further elucidate the animal methods bias phenomenon.

The survey asked a series of questions aimed at understanding the experiences and perceptions of both authors and reviewers during the manuscript submission process. A total of 21 respondents said they have carried out animal-based experiments for the sole purpose of anticipating reviewer requests for them (Tab. 2). A total of 31 respondents said they have been asked for animal experimental data be added to a study that otherwise had no animal-based experiments at least once (Fig. 1). A total of 15 respondents said, in their capacity as reviewers, have made such requests, several indicating that they did so for reasons including “Preference for animal-based methods,” and being “Unaware of nonanimal or alternative methods for the given hypothesis” (Fig. 2). The data presented here supports the occurrence of animal methods bias in publishing, which represents a novel type of bias in scientific publishing.

In a broader sense, bias in scientific publishing can be defined as “a systematic prejudice that prevents the accurate and objective interpretation of scientific studies” (10). According to a handbook on publication bias by the Oxford Handbooks Online, “the importance of addressing publication bias is as fundamental as the importance of promoting good science in general” (11). Bias in the scientific literature may be of different types. Mostly, publication bias is recognized as any practice that distorts the true value of scientific results, producing misleading conclusions that are based on incomplete records of reporting results (only positive results are published) or on results that imply associations, cause-effect relations, prevalence, and other measures that are not supported by the available data or cannot be reproduced by different research groups. There are some suggestions of methodologies to identify these types of biases in the scientific literature. One idea proposed by Ioannidis and Trikalinos in 2007 is based on an exploratory test that evaluates whether the number of studies presenting statistically significant findings available in the scientific literature is higher than expected (12); another idea is to investigate ethical review boards, data repositories, and registries to find information on studies planned and/or intended to be carried out and check whether these have ever been published and how they were reported (13).

The Catalogue of Bias lists more than 50 distinct types of biases in clinical research and health-related studies, yet none describe the bias we report here (www.catalogofbias.org). The type of publication bias we investigated in this study is different than what is commonly recognized as publication bias, and it is not expected to be associated with missing data or findings that are never submitted or accepted for publication due to the quality of the finding itself (mostly negative findings) (1). Instead, it is related to the preference for certain methods to obtain specific data when there may be no scientific justification for such preference. Thus, animal methods bias may be described as a systemic and internal bias within the scientific community, which demonstrates an inherent problem within the culture of science. One major issue stemming from this bias may be a misattribution of success in biomedical advances due to the use of animal-based experiments, despite the foundational work being done in nonanimal human-specific experimental systems. For example, the physiological breathing motion and not the presence of immune cells is the main reason cancer patients develop edema when treated with IL-2, a finding obtained using the lung alveolus chip (14). More recently, the lung alveolus chip helped to elucidate how bacterial lipopolysaccharide endotoxin stimulates intravascular thrombosis (15). Currently, this chip is being used to study infection by SARS-CoV-2 and to test FDA-approved drugs as potential candidates for treating COVID-19 (16). There is no scientific basis to justify a request for animal-based experiments to be performed to “validate” these types of findings obtained with technologies that did not rely on animals, and yet, it has been reported that such requests have been made (4). Because there is no reason to expect that a human-based experimental system would produce the same results as an animal experiment, these requests skew expectations, confuse the scientific literature, and may even increase the use of animals in research.

Animal methods bias in publishing aligns with the concept of conservatism bias, which is bias against groundbreaking and innovative research. In a presentation done at the First International Congress on Peer Review in Biomedical Publication that took place in Chicago in 1989, David Frederick Horrobin warned the audience about how peer review, and more specifically conservatism bias, may hinder scientific progress. In his speech, Horrobin listed around 20 cases of innovations that faced non-scientifically justified resistance and that included the paper describing the tricarboxylic acid cycle, a discovery that honored Hans Krebs a Nobel Prize in 1953 (17). In fact, some of the most cited papers ever did not pass peer review at first, and some took years to be accepted (18). Based on survey respondents’ elaborations (Boxes 1, 4 and 5), animal methods bias may also occur during the peer review of grant applications, which may likewise delay scientific progress. Types of unconscious bias are recognized by biomedical grant-making institutions like the U.S. National Institutes of Health and can take the form of “having more enthusiasm for applications addressing someone’s own area of research,” such as the use of animal-based experimental methods (19).

In our study, it was revealing that, when acting as reviewers, respondents described the influence of editor requests, personal preference to animal-based methods, or lack of awareness of nonanimal methods. These results are in line with a study conducted to investigate peer reviewers’ biases in accepting results produced while using methods not considered as conventional. In this study, authors found that reviewers tend to reject studies done using practices they consider as unconventional, which is supported by other studies showing that reviewers’ judgment of the quality of a scientific study relies on their prior beliefs (20). In fact, a recent study that aimed at investigating how a given animal model is chosen for studying a specific human condition found that the selection of an animal model is based mainly on availability, while the probability that results obtained using the model will translate to humans is not a criterion for choosing the model (21). In 2013, a study comparing gene expression levels and genomic response to inflammation in humans who suffered trauma, burn, or endotoxemia and corresponding mouse models for these conditions received many critiques and was rejected for publication by a number of journals because it presented evidence of virtually no correlation between humans and the animal counterparts for the conditions investigated. Studies that contradict long-term beliefs may also find resistance in the scientific community, and difficulties to be accepted for publication, regardless of how solid the scientific evidence is (22).

In our study, authors who indicated that they had been requested by reviewers to add animal-based experiments to a study that otherwise had no animal-based experiments were asked if they thought the requests were justified. Only three responded yes, while 14 responded sometimes and 11 responded no. When asked to elaborate on this question, eight respondents expressed that the requested experiment would not add value to the study or implied that the request was made out of habit, because editors and reviewers are more familiar with animal data (Box 1). Indeed, the preference for long-time conventions may interfere with people’s judgment when data indicate otherwise. This lack of flexibility may put scientists that work exclusively with nonanimal methods at a disadvantage in publishing. Moreover, studies conducted completely without the use of animals may be underrepresented in journals, creating a false impression that findings from these types of studies are unable to be translated to clinical trials without inclusion of animal data. This convention may delay scientific progress and the full acceptance of technologies that do not rely on animal data. Animal methods bias also adds to the number of background papers that have produced *de novo* animal-data to “validate” findings originally produced with technologies that do not rely on animals, giving the impression that findings obtained with technologies such as microphysiological systems, which better mimic human biology, require additional evidence that can only be provided by animal data. It should be noted that animal data, and more specifically animals, often must simply mimic certain human biological factors, such as gene expression, to be considered a valid experimental model.

Although the current study does not comprehensively explore the consequences of animal methods bias, its impact may be significant and may contribute to the continued use of animal-based research methods, the stifled use and further development of nonanimal methods, and the misdirected prioritization of federal research dollars. Thus, strategies to combat this bias are needed, even as researchers seek to further understand the causal factors of its existence. Some ways to address animal methods bias in publishing may be peer review training and accreditation and a two-stage review. A study that explored practical solutions for reducing publication bias reported that editors, reviewers, and researchers differ on what they see as a solution for the problem of publication bias but most recognized it as a problem associated with peer review (23). In that study, one of the suggested solutions for reducing publication bias was peer review training and accreditation, even though the option was not well accepted by editors in general. Interestingly, the study shows that this option is especially favored by early-career scientists (< 10 years of experience), indicating that these individuals may see the need for training to be prepared for reviewing the work of their peers. The two-stage review process, another of the suggested solutions in which editors and reviewers decide whether to recommend a paper for publication based on the article’s introduction and methods sections only, was considered as the best option for reducing publication bias. The rationale behind this idea is that the decision regarding publication should not depend on the study’s results but on its design and the methods it used. It would therefore be important to investigate how the two-stage review process could address animal methods bias in publishing, as this bias seems to stem primarily from the methods used rather than the results obtained.

The practice of open peer review endeavors to improve the process of peer review by increasing transparency, mirroring the goals of the broader open science movement (24). With open peer review, aspects of the peer review process are made public, such as the name the reviewers assigned to a paper and reviewer comments, including any requests for adding animal data. A 2000 randomized controlled trial of open peer review found that open reviews were of higher quality, were more courteous, and took longer to complete than closed reviews (25). In a 2015 survey of editors aiming to assess the impact of an open review pilot, 33% of editors identified an improvement in the overall quality of review reports, and 70% of those editors said that the pilot resulted in reports that are “more in depth and constructive for authors to improve the quality of their manuscript” (26). Transparency is a crucial component of rigorous science, especially with regard to animal research, a notion upheld by the U.S. National Institutes of Health, the largest funder of biomedical research in the world (27). Open peer review may therefore be a useful tool in addressing animal methods bias by encouraging reviewers to take more time to produce more thoughtful and higher quality reviews.

In addition, open peer review would make it known to readers when experimental results were obtained at the request of reviewers as opposed to being conceived as a part of the original study plan. The “validation” of human-specific models with animal-based data often lacks scientific justification and ethics principles require its avoidance. It is unlikely that performing *de novo* animal experiments to generate data to be added to a study originally conceived without the use of animals can adhere to the 3Rs principle; depending on the country or region, ethical committees may not (or should not) approve uses of animals that were not part of the original study plan.

Currently, the causal factors leading to animal methods bias in publishing are not fully understood. It is possible that in some cases reviewers and editors are unfamiliar with the strengths of nonanimal methods in providing mechanistic knowledge and testing novel therapeutics. As described above, this may result in peer review requests or submission requirements for more conventional, animal-based methods to be used to produce data to “validate’’ scientific findings previously obtained using animal-free methods. Other explanations behind animal methods bias in publishing may be: (i) the occurrence of *status quo* bias, defined as a preference for the current state of affairs, and which is based on emotional rather than objective reasons; (ii) the policy some journals have that findings should be validated *in vivo*, contributing to the perpetuation of animal methods bias in publishing; and (iii) regulatory testing requirements, which may also suffer from this type of bias.

When evaluating the occurrence of any type of bias, the possibility that conflicts of interest (CoI) are at the root of the problem or may contribute to its perpetuation cannot be ruled out. Specifically, financial and professional relationships with animal-based companies or advocacy groups promoting animal-based experiments can increase peer reviewers’ or editors’ internal animal methods bias. Thus, it should be noted that while authors are virtually always asked to disclose any CoI they may have when submitting a paper for publication, the same request does not always apply to reviewers or editors. This situation is observed even in journals that subscribe to professional organizations’ rules such as The Committee on Publication Ethics and the International Committee of Medical Journal Editors that do recommend that both reviewers and editors disclose any conflict of interest they may have. Requiring CoI disclosure from all those involved in peer review, and not only authors, may help address animal methods bias in publishing (28).

A different type of conflict may also take place with animal ethics committees, such as Institutional Animal Care and Use Committees (IACUC). Many institutions have issued rules in order to avoid CoI involving IACUC members. The IACUC is a central organization that oversees animal-based research in the United States and is responsible for ensuring that research involving animals is done according to established regulations. However, no studies or reports describe what happens when scientists request approval by the IACUC to perform an experiment that was requested by reviewers during peer review, and which may be conditional for paper acceptance in a journal. The pressure to publish and the interest of the research group in having the paper accepted for publication may create a CoI for the research group, a situation that warrants further investigation. It may also create a pressured situation for the IACUC to expedite decisions that will favor the research group and the interests of the institution. An avenue for further investigation may be to determine whether the IACUC’s decision to approve an animal experiment is different when the request for approval occurs before manuscript submission and peer review as opposed to when it occurs during review as a potential condition for manuscript acceptance.

This is the first study, to our knowledge, to describe evidence of animal methods bias in publishing and provides the basis for developing strategies to minimize its effects and address some of its potential causal factors. Additional work is needed to further characterize animal methods bias, evaluate how pervasive it is within the many disciplines of science, and identify what role it may play beyond publishing, especially in the review of grant applications. Elimination of this type of bias will ultimately improve science communication and transparency and will help advance the development and adoption of methods in biomedical research that do not rely on animals, are more human-relevant, and aid in translation to the clinic.

## Data availability statement

Survey data is available and can be found in the Supplementary Material (S3 Data).

## Acknowledgments

Thank you to Marcia Triunfol for her contributions to the survey and manuscript. Thank you to Lindsay Marshall and Bianca Marigliani for their remarks and suggestions on how to improve the manuscript. And thank you to the survey respondents for taking the time to share their experiences with us.

## Supplementary Tables and Figures

**Table S1.** Additional respondent demographics.

| Characteristic | N | Percent total |
| --- | --- | --- |
| *Total* | 68 |  |
| Field (question 1) |  |  |
| Anthropology | 1 | 1% |
| Biochemistry and biophysics | 4 | 6% |
| Biotechnology | 8 | 12% |
| Computational biology | 4 | 6% |
| Dentistry | 1 | 1% |
| Disease research | 8 | 12% |
| Ecology | 1 | 1% |
| Environmental sciences | 0 | 0% |
| Evolutionary biology | 3 | 4% |
| Genetics and genomics | 6 | 9% |
| Health disparities | 1 | 1% |
| Medicine and clinical research | 19 | 28% |
| Microbiology | 3 | 4% |
| Molecular and cellular biology | 14 | 21% |
| Neuroscience | 10 | 15% |
| Nutritional sciences | 3 | 4% |
| Pharmacology | 11 | 16% |
| Physiology | 3 | 4% |
| Social sciences | 3 | 4% |
| Systems biology | 2 | 3% |
| Toxicology | 13 | 19% |
| Other | 6 | 9% |
| Country (question 4) |  |  |
| Afghanistan | 1 | 1% |
| Belgium | 1 | 1% |
| Brazil | 5 | 7% |
| Canada | 3 | 4% |
| China | 1 | 1% |
| Denmark | 2 | 3% |
| Ecuador | 1 | 1% |
| Faroe Islands | 1 | 1% |
| Germany | 1 | 1% |
| Greece | 2 | 3% |
| India | 2 | 3% |
| Iraq | 1 | 1% |
| Italy | 4 | 6% |
| Korea, South | 10 | 15% |
| Netherlands | 2 | 3% |
| Pakistan | 1 | 1% |
| Poland | 1 | 1% |
| Romania | 1 | 1% |
| Slovakia | 1 | 1% |
| Switzerland | 1 | 1% |
| United Kingdom | 4 | 6% |
| United States | 22 | 32% |

**Fig S2.**
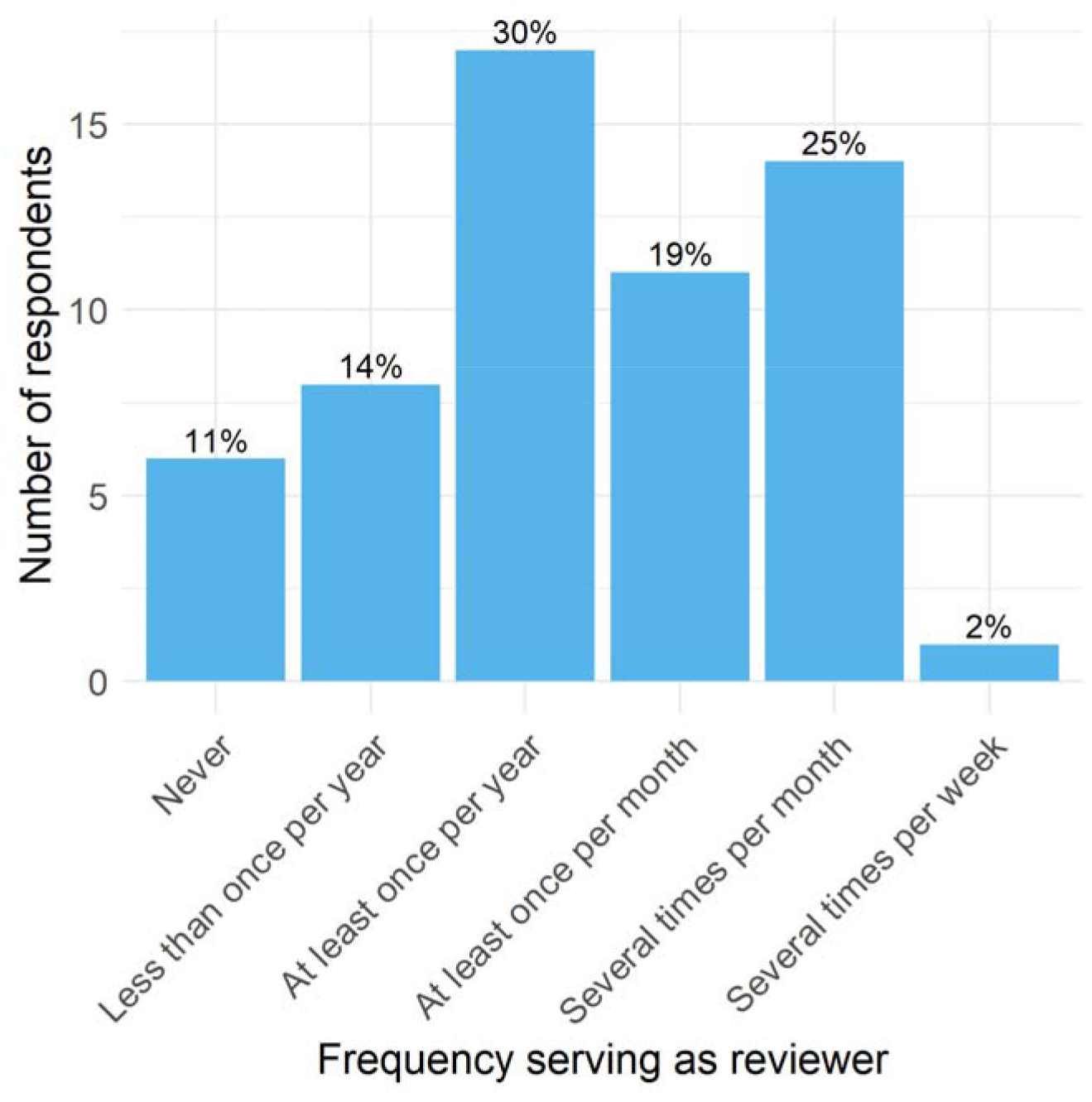
Frequency at which respondents serve as reviewer. Responses to question 25. The percent of respondents answering in each category is on top of each bar (total N= 57).

