## Supplementary material for "A survey to assess animal methods bias in scientific publishing": S1 Appendix

* 1. What best describes your field? Select all that apply.

- [ ] Anthropology
- [ ] Biochemistry and biophysics
- [ ] Biotechnology
- [ ] Computational biology
- [ ] Disease research
- [ ] Ecology
- [ ] Environmental sciences
- [ ] Evolutionary biology
- [ ] Genetics and genomics
- [ ] Health disparities
- [ ] Medicine and clinical research
- [ ] Microbiology
- [ ] Molecular and cellular biology
- [ ] Neuroscience
- [ ] Nutritional sciences
- [ ] Pharmacology
- [ ] Physiology
- [ ] Social sciences
- [ ] Systems biology
- [ ] Toxicology
- [ ] Other (please specify)

* 2. In which sector do you work?

- [ ] Academia/Research institution
- [ ] Industry
- [ ] Government
- [ ] Nonprofit/NGO
- [ ] Publishing
- [ ] Other (please specify)

3. What is your gender?

- Male
- Female
- Nonbinary
- Prefer not to say
- Prefer to self-describe

* 4. In what country do you primarily work?

* 5. What is your highest earned degree?

- Master's degree
- Research doctorate (e.g. PhD)
- Medical degree (e.g. MD, DDS)
- Other (please specify)

* 6. How many years has it been since you received your highest earned degree?

- 0-5
- 6-10
- 11-20
- >20

* 7. How many peer-reviewed publications do you have?

- 0
- 1-10
- 11-20
- 21-40
- 41-100
- >100

* 8. Do you regularly use animal or nonanimal models, samples, data, or materials in your research? Select all that apply.

| | Models | Samples | Data | Materials | None |
|---|---|---|---|---|---|
| Animal-based | ☐ | ☐ | ☐ | ☐ | ☐ |
| Nonanimal-based | ☐ | ☐ | ☐ | ☐ | ☐ |
| Other (please specify below) | ☐ | ☐ | ☐ | ☐ | ☐ |

Other (please specify)

* 9. What kind of model(s) do you regularly use in your research? Select all that apply.

- ☐ 2D cell culture
- ☐ Biopsies
- ☐ Computational
- ☐ Live organisms
- ☐ Microphysiological systems, organ chips, tissue chips
- ☐ Organoids, spheroids
- ☐ Post-mortem tissue
- ☐ Other (please specify)

* 10. How often do you perform animal-based experiments?

- ○ Never
- ○ Rarely
- ○ Sometimes
- ○ Often
- ○ Always

* 11. How often do you perform animal-based experiments for the sole purpose of anticipating reviewer requests for them (i.e., you did not think the experiments were necessary outside of the context of review)?

- Never
- Rarely
- Sometimes
- Often
- Always

* 12. During manuscript submission peer review, how many times have you been asked for animal experimental data to be added to a study that otherwise had no animal-based experiments?

- 0
- 1-5
- 6-10
- 11-20
- >20
- N/A; e.g. if you've only submitted studies with animal-based experiments

* 13. Did you feel the requested additional animal-based experiments were justified?

- Yes
- No
- Sometimes
- Not sure

14. Please elaborate on your answer to the previous question.

* 15. Did you comply with the request(s)?

- Yes
- No
- Sometimes

16. Please elaborate on your answer to the previous question.

* 17. When you complied with the request(s), did members of your own lab or the labs of coauthors perform additional experiments, or did you and coauthors bring in additional collaborators?

- Own lab(s) or lab(s) of coauthors
- Additional collaborators
- Combination
- Other (please specify)

18. Please elaborate on your answer to the previous question.

* 19. Have you ever had a manuscript rejected because you did not comply with a request for animal experimental data to be added to a study that otherwise had no animal-based experiments?

- Yes
- No
- Sometimes
- Not sure

20. Please elaborate on your response to the previous question.

* 21. During manuscript submission peer review, how often have you been asked for additional nonanimal-based experimental data to be added to a study that had no animal-based experiments? *

- Never
- Rarely
- Sometimes
- Often
- Always
- N/A

22. During manuscript submission peer review, how often have you been asked for nonanimal-based experimental data to be added to a study that had animal-based experiments? *

- Never
- Rarely
- Sometimes
- Often
- Always
- N/A

* 23. During manuscript submission peer review, how often have you been asked for additional animal-based experimental data to be added to a study that had animal-based experiments?

- Never
- Rarely
- Sometimes
- Often
- Always
- N/A

24. Are there any additional details you would like to share about your experience(s) during manuscript review being asked for additional experimental data?

25. How often do you serve as a manuscript reviewer for scientific journals? *

- Never
- Less than once per year
- At least once per year
- At least once per month
- Several times per month
- Several times per week

* 26. In your capacity as a reviewer for a scientific journal, how often do you request animal experimental data to be added to a study that otherwise had no animal-based experiments?

- ◯ Never
- ◯ Less than once per year
- ◯ At least once per year
- ◯ At least once per month
- ◯ Several times per month
- ◯ Several times per week
- ◯ N/A

* 27. For what reasons have you requested that animal-based experimental data be added to a study that otherwise had no animal experiments?

- ☐ Journal policy
- ☐ Request from editor
- ☐ Preference for animal-based methods
- ☐ Unaware of nonanimal or alternative methods for the given hypothesis
- ☐ Other (please specify)

28. Please elaborate on your response to the previous question.

* 29. In your capacity as a reviewer for a scientific journal, how often do you request nonanimal-based experimental data to be added to a study that had animal-based experiments?

- Never
- Less than once per year
- At least once per year
- At least once per month
- Several times per month
- Several times per week
- N/A

* 30. In your capacity as a reviewer for a scientific journal, how often do you request nonanimal-based experimental data to be added to a study that had no animal-based experiments?

- Never
- Less than once per year
- At least once per year
- At least once per month
- Several times per month
- Several times per week
- N/A

* 31. In your capacity as a reviewer for a scientific journal, how often do you request animal-based experimental data to be added to a study that had animal-based experiments?

- Never
- Less than once per year
- At least once per year
- At least once per month
- Several times per month
- Several times per week
- N/A

32. Are there any additional details you would like to share about your experience(s) as a reviewer requesting additional data?

33. The authors of this survey would greatly appreciate the opportunity to discuss your responses in greater detail to gather additional qualitative data regarding reviewer requests for additional animal evidence. Your further participation will remain entirely confidential. If you are interested, please provide contact information.

| |
|---|
| **Name** |
| **Email Address** |
| **Phone Number** |
