## Supplementary material for "A survey to assess animal methods bias in scientific publishing": S2 Appendix

**S3 Appendix: Survey logic**

Question 1: No logic

Question 2: No logic

Question 3: No logic

Question 4: No logic

Question 5: No logic

Question 6: Skip logic

- Answer 1. 0: Skip to end of survey
- Answer 2. 1-10: No logic
- Answer 3. 11-20: No logic
- Answer 4. 21-40: No logic
- Answer 5. 41-100: No logic
- Answer 6. >100: No logic

Question 7: No logic

Question 8: No logic

Question 9: No logic

Question 10: No logic

Question 11: Skip logic

- Answer 1. 0: Skip to question 20
- Answer 2. 1-5: Skip to question 12
- Answer 3. 6-10: Skip to question 12
- Answer 4. 11-20: Skip to question 12
- Answer 5. >20: Skip to question 12
- Answer 6. N/A: Skip to question 20

Question 12: No logic

Question 13: No logic

Question 14: Skip logic

- Answer 1. Yes: No logic
- Answer 2. No: Skip to question 18
- Answer 3. Sometimes: No logic

Question 15: No logic

Question 16: No logic

Question 17: No logic

Question 18: No logic

Question 19: No logic

Question 20: No logic

Question 21: No logic

Question 22: No logic

Question 23: Skip logic

- Answer 1. Never: Skip to question 29
- Answer 2. Less than once per year: Skip to question 24
- Answer 3. At least once per year: Skip to question 24
- Answer 4. At least once per month: Skip to question 24
- Answer 5. Several times per month: Skip to question 24
- Answer 6. Several times per week: Skip to question 24

Question 24: Skip logic

- Answer 1. Never: Skip to question 27
- Answer 2. Less than once per year: No logic
- Answer 3. At least once per year: No logic
- Answer 4. At least once per month: No logic
- Answer 5. Several times per month: No logic
- Answer 6. Several times per week: No logic

Question 25: No logic

Question 26: No logic

Question 27: No logic

Question 28: No logic

Question 29: No logic
